# Diagrammatic Theory of RNA Structures and Ensembles with Trinucleotide Repeats

**DOI:** 10.1101/2020.05.30.125641

**Authors:** Chi H. Mak, Ethan N. H. Phan

## Abstract

Trinucleotide repeat expansion disorders (TRED) are associated with the overexpansion of (CNG) repeats on the genome. mRNA transcripts of sequences with greater than 60 to 100 (CNG) tandem units have been implicated in TRED pathogenesis. In this paper, we develop a diagrammatic theory to study the structural diversity of these (CNG)_n_ RNA sequences. Representing structural elements on the chain’s conformation by a set of graphs and employing elementary diagrammatic methods, we have formulated a renormalization procedure to resum these graphs and arrive at a closed-form expression for the ensemble partition function. With a simple approximation for the renormalization and applied to extended (CNG)_n_ sequences, this theory can comprehensively capture an infinite set of conformations with any number and any combination of duplexes, hairpins and 2-way junctions. To quantify the diversity of different (CNG)_n_ ensembles, the analytical equations derived from the diagrammatic theory were solved numerically to derive equilibrium estimates for the secondary structural contents of the chains. The results suggest that the structural ensembles of (CNG)_n_ repeat sequence with n ~ 60 are surprisingly diverse, and they are dominated largely by open segments, with only a small fraction of the nucleotides forming base pairs. At the same time, the variance in the secondary-structural contents on the chains is also quite large, indicating that their structures can undergo strong equilibrium fluctuations and are expected to be rather suspectable to perturbations.

**STATEMENT OF SIGNIFICANCE:** Trinucleotide repeat expansion disorders (TRED) are associated with the overexpansion of (CNG) repeats on the genome. mRNA transcripts of sequences with critical length greater than 60 to 100 (CNG) tandem units have been implicated in TRED pathogenesis, though their structures remain poorly characterized. Conventional view has tacitly assumed that conformations with maximal C:G base pairing dominate at equilibrium, but here we demonstrate that (CNG) repeat sequences are characterized by diverse ensembles of structurally heterogeneous folds and with a large variance of secondary structural contents. These ensembles of structures also undergo strong equilibrium fluctuations, rendering them rather susceptible to perturbations. These results were based on a novel diagrammatic approach to the ensemble partition function.

## INTRODUCTION

A graphical approach for classifying RNA structures has been pioneered by Schlick et al. (1–3). Graphs provide an elegant method for categorizing the many diverse conformational structures that can be adopted by RNA sequences and can help us more easily recognize common topological features in RNA structures that are otherwise difficult to decipher from their 2-or 3-dimensional structures. But more than assisting in cataloging RNA motifs, graph can also provide clues to the “factorizability” of different classes of substructures in RNA folds (4) {companion paper}. Here, we develop these ideas further and formulate a diagrammatic theory to take advantage of these factorizabilities, applying it to evaluate the partition functions of RNA structure ensembles and to study the conformational diversities in their folded structures.

To appreciate what these factorizabilities means for RNA structures, consider an open RNA strand. An unfolded RNA is in a high-entropy state. Its structures are characterized by a diverse ensemble. If *c* denotes a chain conformation and *P*(*c*) its probability, the total entropy content of this ensemble is given by 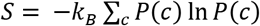. If the sequence spontaneously folds and develops secondary and/or tertiary structures, the conformational entropy of the chain is suppressed because base complementarity and stacking interactions produce constraints on the chain’s conformations. Under these constraints, the new probability for each conformation in the presence of these constraints *P*′(*c*) = *P*(*c*|constaints) incurs a penalty, and the loss of entropy is given by:

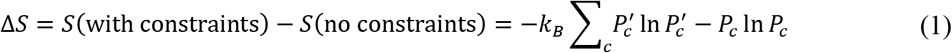

where the sum runs over all conformations. If one can determine how the constraints imposed by the secondary and tertiary structures in the fold transforms *P*(*c*) → *P*′(*c*), Δ*S* can be determined.

In general, the constraints imposed by secondary/tertiary structures are correlated, but some of them may be independent. “Factorizability” describes how these constraints break up into independent (or approximately independent) subsets. For instance, if the fold introduces 4 constraints *A*, *B*, *C* and *D* but the effects of *A* and *B* produce are separable from *C* which is also separable from *D*, then *P*′(*c*) = *P*(*c*|*A*, *B*, *C*, *D*) = *P*(*c*|*A*, *B*) ⋅ *P*(*c*|*C*) ⋅ *P*(*c*|*D*). Under this factorization, the entropy change in Eq.(1) is simply equal to Δ*S* = Δ*S*(with constraints *A*, *B*) + Δ*S*(with constraint *C*) + Δ*S*(with constraint *D*).

In Schlick’s graph notations, the secondary structures inherent in a RNA fold are represented by edges and vertices (1–3). Using a large body of empirical data derived from Monte Carlo conformational sampling (4) {companion paper}, we have derived a set of general rules for determining which constraints are independent and how they can be divided into uncorrelated subsets. When attempting to summarize these rules visually, we discovered that there is a close correspondence between factorizability and how these constraints are depicted in the graphs proposed by Schlick. By devising an appropriate set of rules, these diagrams can be used to easily factorize the constraints. Once factorized, the diagrams can then be used to represent the partition function of the entire ensemble. In this paper, we illustrate how graphs can also be used to construct resummation techniques to sum the diagrams, leading to an analytical expression for the partition function. We will apply this technique to examine the diversity of structures in ensembles of (CNG)_n_ trinucleotide repeat RNA sequences.

In a diverse family of neurological diseases known as trinucleotide repeats expansion disorders (TREDs) (5–9), the onset of illness is associated with the overexpansion of (CNG)_n_ repeats in the genome (8–10). While most of these expanded repeats occur in noncoding regions and do not appear to translate to aberrant proteins (9, 10), the mRNA transcripts of these overexpanded templates may interfere with cellular pathways leading to cytotoxicity(11, 12). At the same time, (CNG)_n_ expanded mRNA may also acquire unintended functions in the cell (13). Ascertaining the structures of these sequences is therefore necessary for the understanding of their functions.

Two examples of a short (CNG) repeat with different secondary structures are shown in Fig. 1. Because of their repeat structures, at least one-third of the nucleotides on (CNG)_n_ sequences cannot form complementary pairs upon folding. TRED disease onset is often associated with a critical expansion threshold of n > 60 to 100 (14). The structures most often associated with the gain of function hypothesis for CNG expanded RNA sequences cited in the literature is a necklace-like structure composed of a long stretch of successive two-way junctions interposed by shorts helixes and with a hairpin stem-loop cap (10, 11, 15–19) like that shown in Fig. 1(a). Many of the studies conducted are based on short (CNG) repeats (10, 11) and the structures resolved are limited to those which can be isolated and crystalized (17–20). As the length of the CNG repeats grows, the diversity of accessible structures could grow rapidly as well. In this paper, we use the diagrammatic theory developed here to study the structural ensembles of long (CNG) repeats.

**Figure 1.**
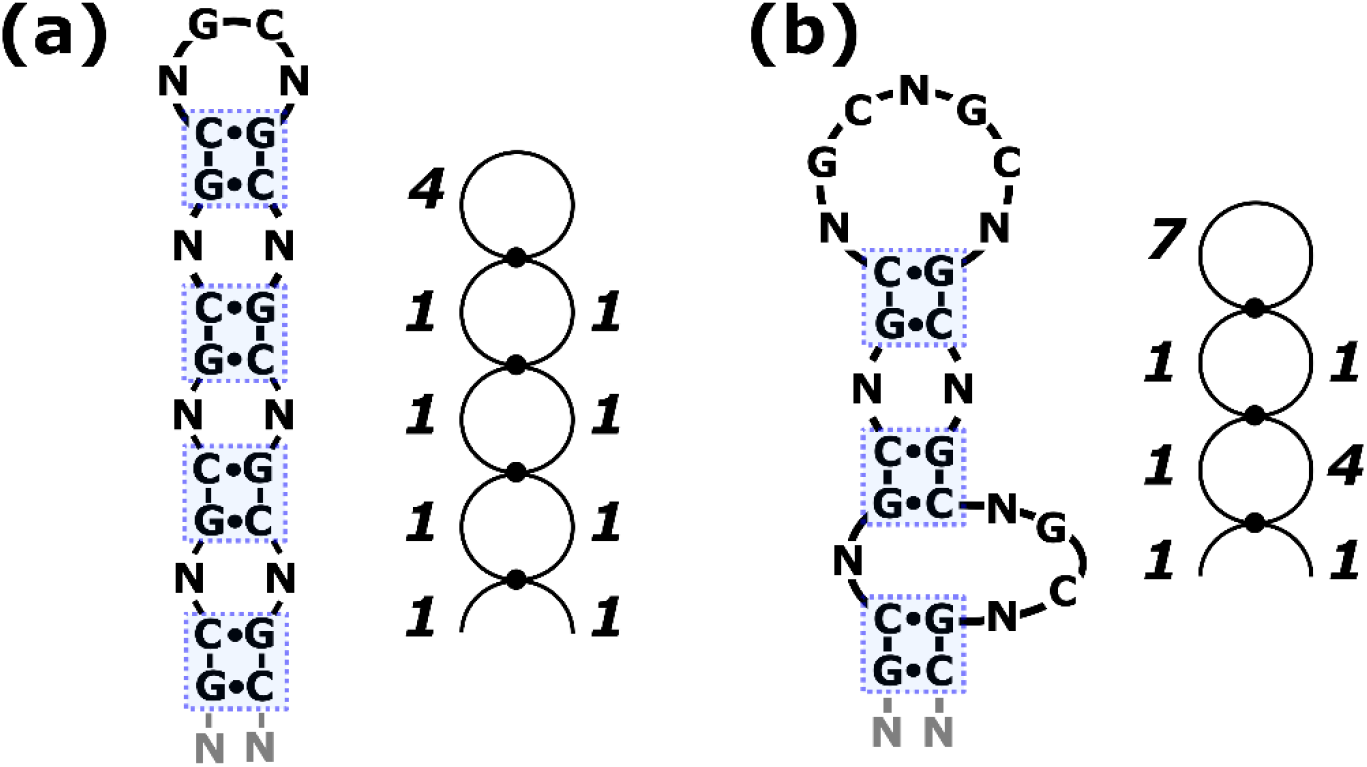
Examples of a (CNG)_9_ repeat sequence in two different conformations. The graph representation for each is shown next to it. Each 2-bp duplex is represented by a dot. A hairpin loop is represented by a circle with one dot. A 2-way junction is represented by a circle with two dots. An arc represents two dangling ends. In the graphs, the number adjacent to each edge indicates its length in nt.

The methods described in this paper are applicable to RNA sequences with many transposable repeats. In this context, “transposable” does not mean mobile. Instead, transposable means that if a segment on the sequence is moved to another position, it does not alter the nucleotide sequence. In the formulation below, it will become clear why transposability is an inherent feature of diagrammatic theories. But a chain with a specific nucleotide sequence cannot form base complementarity interactions equally well among every pair of nucleotides across the entire sequence. As a result, the constraints produced by the secondary structures also have an additional sequence-position dependence, and this renders non-repeat sequences non-transposable. The methods developed here are suitable for treating homopolymeric chains or chains with a large number of regular repeats, and the techniques described in this paper have been formulated with these systems in mind. In the future, we will extend this method to more complex, less regular, and more diverse nucleotide sequences.

## MATERIALS AND METHODS

### Graph Representations

Fig. 2 shows examples of the basic graph elements. The top row shows a 3-way junction in (a), a pseudoknot in (b), a triplex in (c) and a quadruplex in (d). The bottom row shows graph representation of each. Paired regions are represented by dots, triplexes by triangles and quadruplexes by squares. These secondary structures impose constraints on the chains and, as a consequence, produce a loss of conformational freedom in the loop regions. The graphs have been designed to emphasize these features. The middle row in Fig. 2 illustrates pictorially how to transform each of the 2-dimensional drawing in the top row to arrive at the corresponding graph on the bottom row.

**Figure 2.**
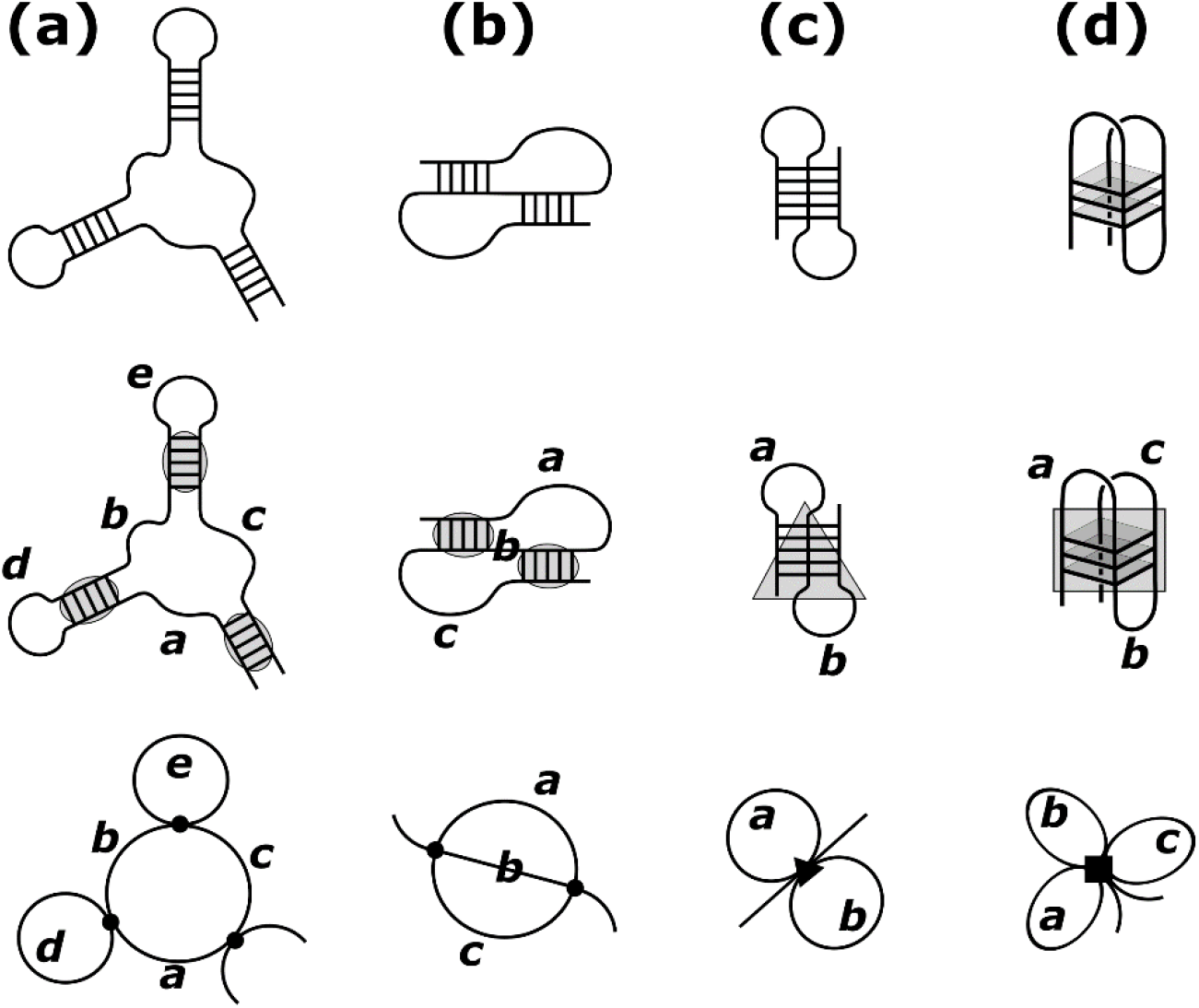
Examples of diagrams representing different secondary structural elements. (a) A three-way junction where each dot represents a helix and loops are represented by lines. (b) A pseudoknot where black dots represent paired regions and unpaired loops are represented by lines. (c) A triplex structure represented diagrammatically by a black triangle. (d) A quadruplex represented by a black square.

We have demonstrated in (4) that most helices are “insulating”, i.e. the two elements immediately adjacent to a helix are approximately uncorrelated. With Fig. 2(a) as an example, the helix on the left and the one on top emanating from the 3-way junction in the center render the two hairpin loops largely uncorrelated with the three unpaired strands on the junction. The open arcs representing the 5’ and 3’ ends of the sequence experience no constraints, and they therefore cost no entropy. In the companion paper {companion paper}, we provide additional data demonstrating the factorizability of pseudoknots and quadruplexes. The factorizability conditions for pseudoknots and quadruplexes are more complex, but they will not be considered in this paper.

Utilizing these graph elements, Fig. 1(a) and (b) show diagrammatic representations for two different conformations of a (CNG)_8_ repeat sequence. The graph associated with Fig. 1(a) shows that it has one hairpin loop and three 2-way junctions. The graph for Fig. 1(b) has one hairpin loop and two 2-way junctions. The loop size in nt is indicated next to each segment. All duplexes in (CNG) repeat structures are 2 bp (4 nt) in length, and since they are uniformly identical for all (CNG) repeat sequences, their sizes are not explicitly labeled in Fig. 1. These graphs resemble pearl necklaces; we will therefore refer to them as “necklace” diagrams. These necklace conformations are the ones generally thought to be most relevant for (CNG) repeat sequences in TREDs (10, 15).

Applying the factorization scheme to the structures in Fig. 1, each graph can be written as a product of independent probabilities. For example, the factorization of the graph in Fig. 1(b) is illustrated in Fig. 3, where the necklace separates into one hairpin loop of length *d*, two 2-way junctions, one with loop lengths *b* and *f* and the other with *c* and *e*, two dangling ends with lengths *a* and *g*, plus three 2-bp (or 4-nt) duplexes. The composite probability of the graph on the left of Fig. 3 is therefore given by:

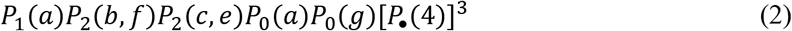

where *P*_1_(*x*) is the probability associated with a hairpin loop (or a “1-way junction”) of length *x*, *P*_2_(*x*, *y*) is the probability of a 2-way junction with loop lengths *x* and *y*, *P*_0_(*x*) = 1 is the probability associated with an open strand and *P*_•_ is the probability of the duplex. For the loops in hairpin and junctions, their probabilities are given by *P* = *e*^Δ*S*/*k*_*B*_^, where Δ*S* is the conformational entropy of a loop relative to an open strand. 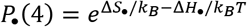, the probability of a 2-bp (4-nt) duplex, has both enthalpic and entropy contributions in it, which involve stacking and base pairing interactions as well as the loss of conformational freedom suffered by the backbone to stack. An example of all the decomposable factors of a necklace diagram is given on the right side of Fig. 3.

**Figure 3.**
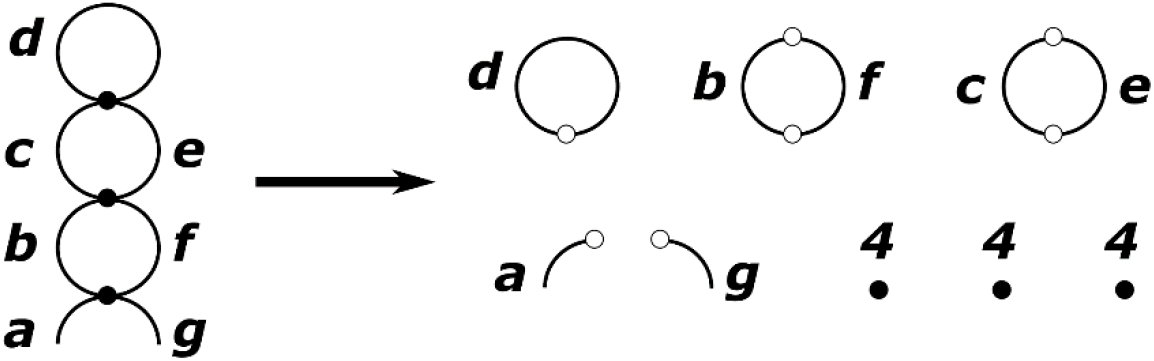
Example showing factorization of the diagram on the left into the factors on the right. The circle with one dot represents a hairpin loop of size *d*. Circles with two dots represent 2-way junctions. The two open line segments represent open strands. The three filled dots represent 2-bp (4-nt) duplexes. The corresponding expression for the composite probability is given in Eq. (2).

### A Crude Approximation for the Partition Function from Necklace Diagrams

Since necklace diagrams are expected to be the ones most relevant to (CNG) repeats, we will use them to illustrate how to calculate the partition function *Z* from the graphs. The complete set of unbranched necklace diagrams is shown in Fig. 4. Let the length of the chain be *l* nt. The sum of all segments in every diagram in Fig. 4 must equal *l*. To sum over all conformations consistent with these graphs, we must trace over all possible combinations of segments subject to the constraint that they sum to *l*. For example, the second term on the right in Fig. 4 corresponds to:

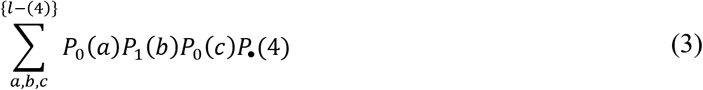

where the sums over the lengths of the three segments *a*, *b* and *c* are constrained such that *a* + *b* + *c* = *l* − 4, which is the length of the chain minus the number of nucleotides in the duplex. Similarly, the third term corresponds to:

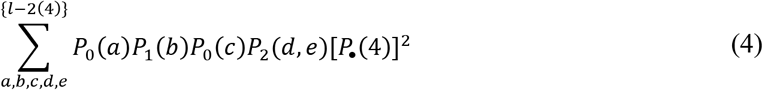

**Figure 4.**
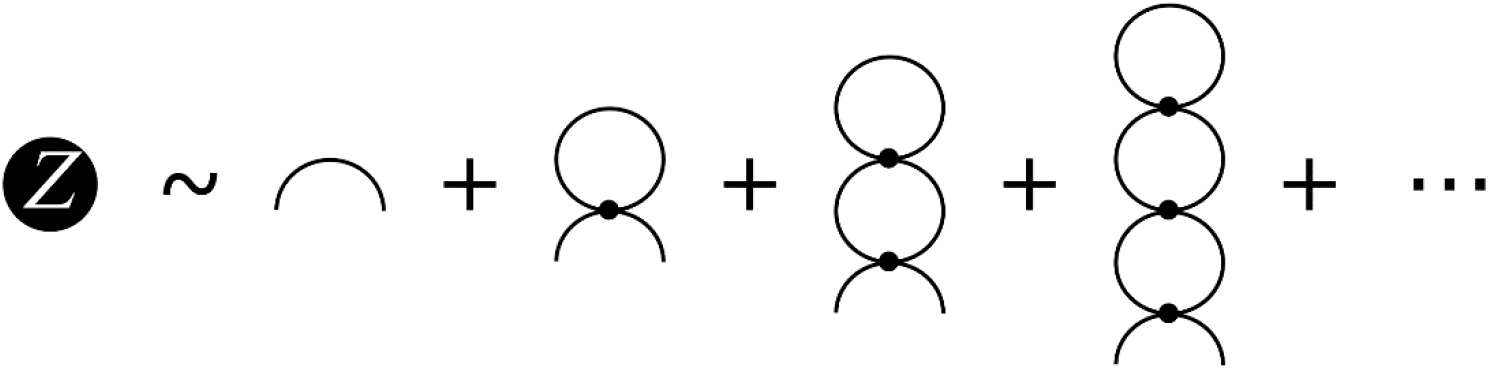
First four terms in an approximation for the partition function *Z* including only nonbranched necklace diagrams for the (CNG) repeats. The corresponding expression is given in Eq.(5).

Writing out the terms explicitly in Fig. 4, the partition function for a *l*-nt chain is:

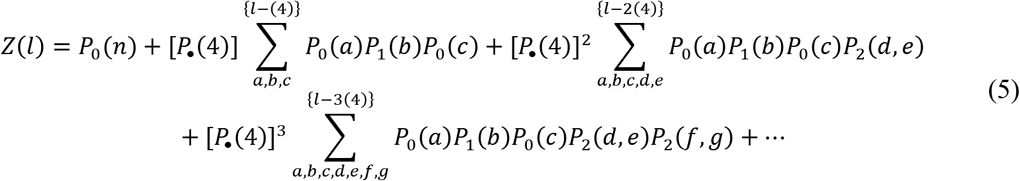

The constrained sums in Eq.(5) are implied in their corresponding diagrams in Fig. 4.

The constraints in Eq.(5) for the partition function are cumbersome. Instead, consider the generating function of *Z*:

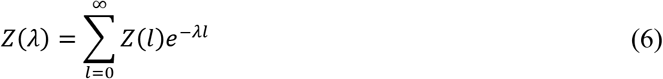

Notice that we have used the same symbol *Z* for the fixed-*l* partition function as well as its generating function. This should not cause any confusion because in the rest of this paper only the generating function *Z*(*λ*) will be considered. The generating function serves the same purpose as the grand canonical partition function, with *λ* corresponding to the chemical potential divided by *k*_*B*_*T*. Using the generating function removes the constraints in *Z* for fixed *l*, and Eq.(5) becomes:

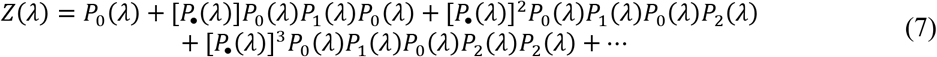

where all the *P* functions are now generating functions of those in Eq.(5). In Eq.(7), the chain length is no longer fixed, but it is possible to compute the expectation of the length ⟨*l*⟩, so that the value of *λ* can be chosen to match the average ⟨*l*⟩ to the proper target chain length *l*. The graph representation for Eq.(7) for the generating functions is identical to that shown in Fig. 4, but the interpretation is distinct.

### Special Graph Elements

We define a few special graph elements in this section in order to facilitate the renormalization scheme in the rest of the paper.

#### a. Dangling Ends and Bridges

The RNA constructs under consideration here always have “dangling ends” on both their 5’ and 3’ ends. Since these dangling ends correspond to open strands, they cost no entropy. Diagrammatically, these correspond to the arcs in Fig. 4. Mathematically, they correspond to *P*_0_(*λ*).

Diagrams can also have “bridges”. A bridge is an edge which if removed from the graph will separate the graph into two disconnected pieces. Examples are shown in Fig. 5. Diagrammatically, bridges also correspond to arcs or non-loop edges, and mathematically, they also correspond to *P*_0_(*λ*). Bridges and dangling ends are topologically equivalent. They cost no entropy and correspond to open strands with probability *P*_0_(*λ*) given by:

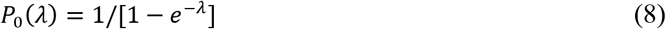

**Figure 5.**
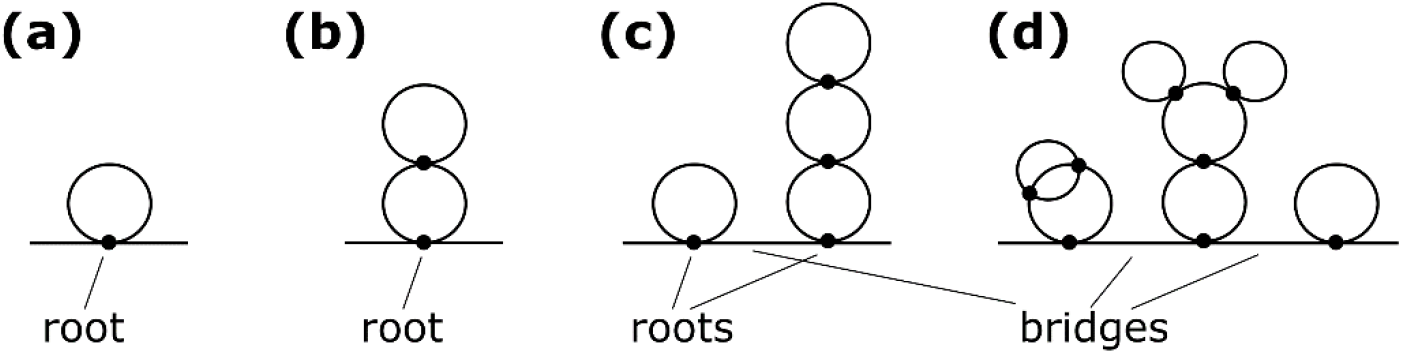
Diagrams showing examples of root vertices and bridge edges.

#### b. Root Vertices

The vertex between two bridges or dangling ends is called a “root”. Examples are shown in Fig. 5. In Ref. 4, we demonstrated that all graphs representing RNA folds are Eulerian and because of this property, the structure of every graph must consist of one or more root vertices connected by bridges, except for the one with just a single edge. Some examples are shown in Fig. 5. A root vertex is simply the beginning and end of a subpart of the diagram that itself forms an Eulerian circuit. Other than the bridges between them, these subparts of the diagram rooted in root vertices do not touch each other.

#### c. Fragile Vertices

Vertices that are defined as “fragile” are the ones which if they are removed from the diagram factor the diagram into disconnected pieces. Examples are the duplexes in Fig. 3 between the hairpin loop and the 2-way junctions, as well the duplexes connect one 2-way junction to the next. As the factorization on the right side of Fig. 3 shows, all helices (dots) in that diagram are fragile.

In contrast, example of vertices that are nonfragile are shown in Fig. 6. These include paired regions in a pseudoknot, as well as base contacts in triplexes and quadruplexes. Empirically, these vertices are found not to factorize completely. For example, the three segments in a pseudoknot are correlated.

**Figure 6.**
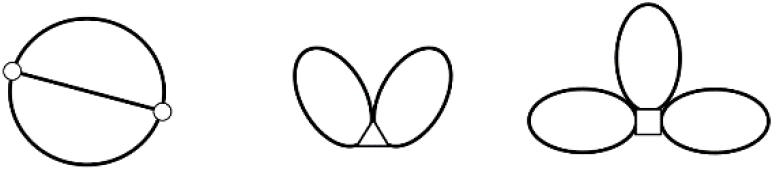
Examples of nonfragile vertices.

Similarly, the two loops in a triplex are also not independent. The three loops inside a quadruplex are also not independent.

#### d. Node Vertices

“Nodes” are topologically similar to roots, except they are not adjacent to bridges. Some examples are shown in Fig. 7 to illustrate how they differ from roots. A node is essentially a branch point. It is the beginning and end of a subpart of the diagram that itself forms an Eulerian circuit, which when the node is removed, would completely disconnect from the rest of the graph.

**Figure 7.**
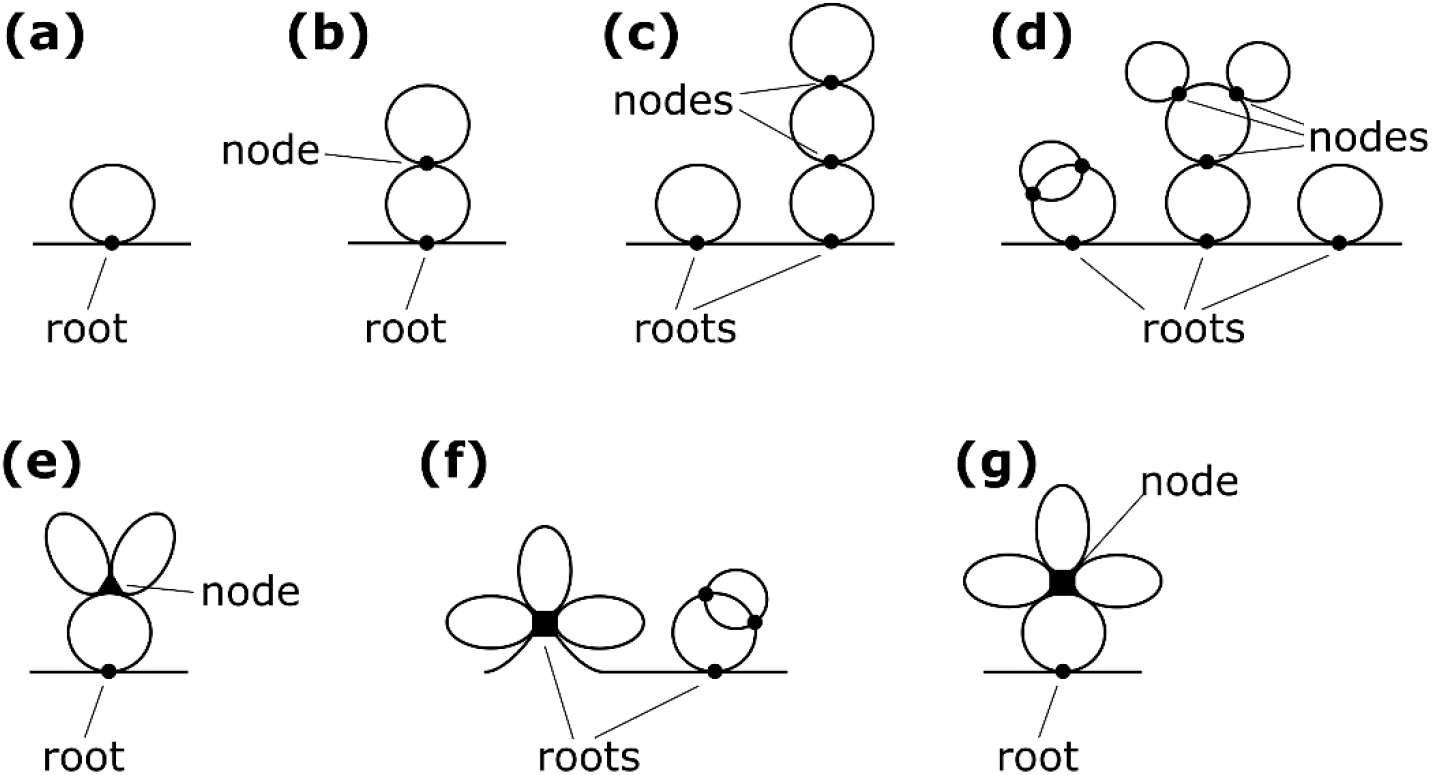
Examples of node vertices and how they differ from root vertices.

### Root Renormalization and Necklace Approximation for the Root Function

The graphs described here share many similar features with those employed in quantum field theory and in liquids, where diagrammatic techniques have been used extensively to manipulate graphs(21). These techniques will be employed here for RNAs.

Root “renomalization” is a diagrammatic reclassification technique. Under root renormalization, all diagrams in the partition function are regrouped into terms forming a simple series shown in the first equality in Fig. 8. Here, the “root function” *R*(*λ*) represents the sum over all possible graphs that begin and end on a root node. Some examples of what are included in the root function *R*(*λ*) are shown in Fig. 9 (which is only a partial list and clearly a very small subset of the full set). How the power series in *R* can be resummed using this regrouping is demonstrated by the second equality in Fig. 8. This yields the result given diagrammatically on the last line in Fig. 8, producing the following *exact* analytical expression for *Z*:

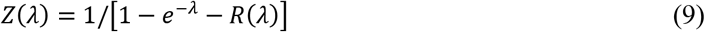

**Figure 8.**
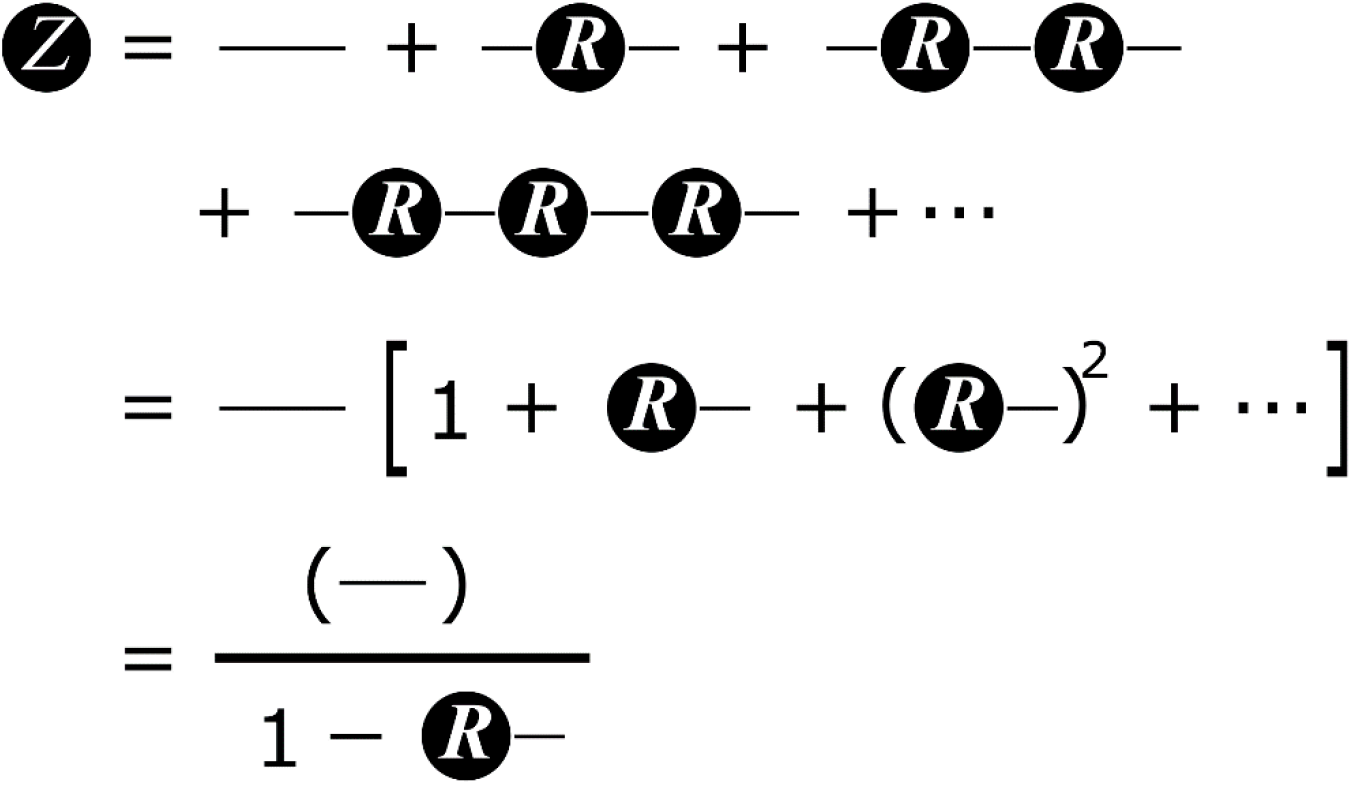
Root renormalization of the partition function *Z*.

**Figure 9.**
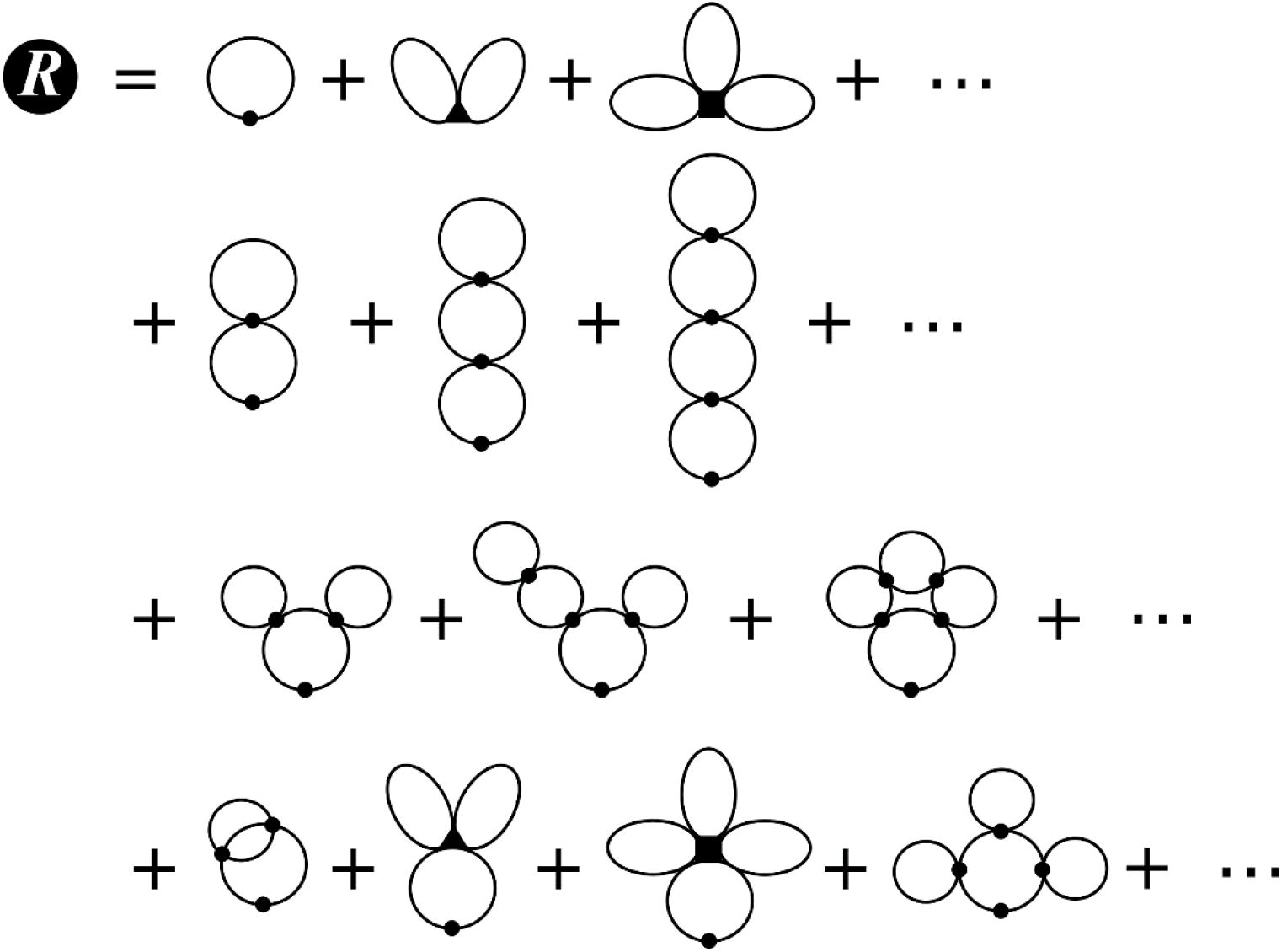
Some examples of the diagrams in the root function *R*(*λ*).

Under root renormalization, the partition function *Z*(*λ*) is re-expressed in a succinct analytical function, in terms of the root function *R*(*λ*). While this seems tantalizing, the root function itself is not known in closed form, so the practical utility of Eq.(9) is limited. To make progress, approximations are often invoked to derive different models for *R*(*λ*). Employing different approximates for *R*(*λ*) in the renormalization of the root then generates distinct classes of theories.

Here, we will consider a “necklace approximation”, *R*_*n*_(*λ*), for the root function. This approximation is shown in Fig. 10. Again, loops with one dot represent hairpins, and those with two dots represent 2-way junctions. Since each 2-way junction interposes between two duplex stems and every circuit beginning and ending on a root vertex must have at least one hairpin, the root function *R*_*n*_(*λ*) includes every diagram with any number of 2-way junctions and a hairpin. Fig. 10 shows how this series is easily resummed, yielding a closed-form expression for the root function:

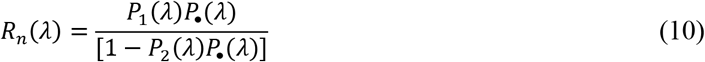

**Figure 10.**
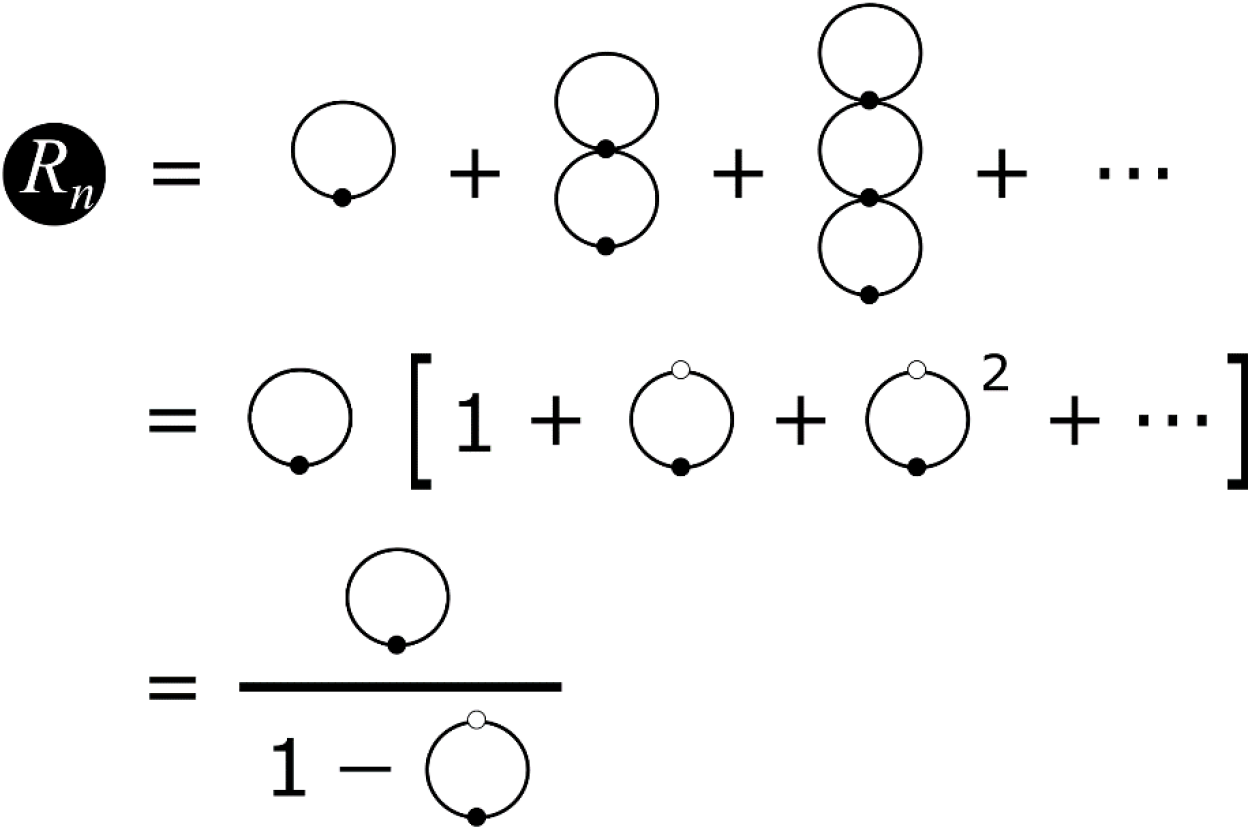
Necklace approximation for the root function and its resummation.

Substituting Eq.(10) into Eq.(9) produces the necklace approximation for the partition function:

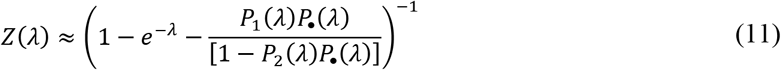

Notice that the partition function *Z*(*λ*) in Eq.(11), obtained by using the necklace approximation to the root function *R*(*λ*), contains *many more diagrams* than the necklace approximation to *Z*(*λ*) in Fig. 4. The much larger set of diagrams included in Eq.(11) contains graphs with any number of roots and bridges connecting them, whereas Fig. 4 is limited to graphs with just one root. Using the necklace approximation *R*_*n*_(*λ*) for the root function, *Z*(*λ*) captures an infinite set of diagrams, including every possible combination of hairpins, 2-way junctions and bridges.

The root function obeys a recurrence relationship that can be derived from a similar renormalization procedure applied to the nodes instead. In a future paper, we will address node renormalization strategies in greater detail. But essentially, the result of this on the root function is to generate an equation of the Dyson type(21), shown in Fig. 11. Solving this equation for *R*_*n*_(*λ*) leads to the same result in Eq.(10). But for cases where the direct resummation of the approximate equation for the root function is not possible, the Dyson equation exemplified by Fig.11 provides an alternate route to the solution.

**Figure 11.**
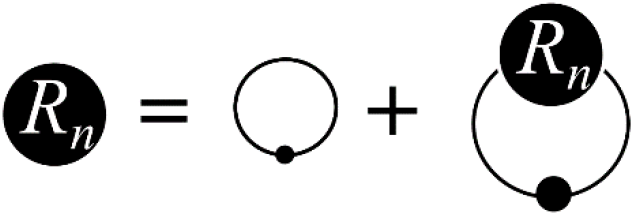
Dyson equation for the root function in the necklace approximation.

### Specializing to (CNG) Repeat Sequences

The theory above has been formulated for generic RNA sequences. To specialize the formulation to apply to (CNG)_n_ repeat sequences specifically, we take a closer look at their repeat structure. By “repeat structure”, we are referring to the periodicity of the nucleotide sequence. In our calculations, we employ (CNG)_n_ constructs with the following architecture:

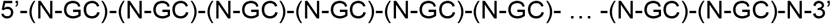

with *n* repeating units of (NGC). Formally, this (CNG)_n_ construct has *l* = 3*n* + 1 nucleotides instead of 3*n*. This is done to ensure that the 5’ and 3’ ends of the chain do not have to be treated differently, but it does not materially alter the results or the formulation.

As described above, the periodicity of the (CNG)_n_ sequence permits canonical base pairing producing 2-bp duplexes only, and we assume the base N does not pair with each other or with C or G (though this assumption can be modified and easily incorporated to re-formulate the theory). Because of this repeat structure, unpaired segments on the sequence are limited to lengths equal to 1, 4, 7, 11,…nt. This means that the sums over *l* used to define the generating functions, for example the one for the partition function in Eq.(6), should be restricted to multiples of 3 instead. Consequently, instead of summing over *l* in Eq.(6), we sum over trinucleotide units *n*. And with just this adaptation, all of the diagrammatic arguments and the equations developed above carry over without any modification, with the proviso that we count in trinucleotide units instead of single nucleotides. To do this, every loop length in the formulation should be replaced by its length divided by 3. For example the lengths {*a*, *b*,…*g*} in Fig. 3 become {*a*′ = *a*\3, *b*′ = *b*\3,…*g*′ = *g*\3}, where \ denotes an integer division without remainder. A loop with length *a*′ = 0 is 1-nt long. A loop with *a*′ = 1 is 4-nt long, etc. The only exception to this rule is a 2-bp (4-nt) duplex, which is assigned a length of 2 repeat units instead of 1.

## RESULTS AND DISCUSSION

The diagrammatic approach formulated here was applied to (CNG)_n_ repeat sequences, using the necklace approximation in Eq.(11) for the generating function of the partition function *Z*(*λ*). The inputs, *P*_•_(*λ*), *P*_1_(*λ*), and *P*_2_(*λ*), were obtained from the free energies of formation of duplexes, hairpins and 2-way junctions from Monte Carlo data reported in an earlier paper (4). The functional dependence of the loop free energies on the loop lengths were extended beyond the finite-length data available from the simulations by using scaling relationships, yielding the following expressions at *T* = 310 K:

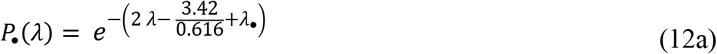

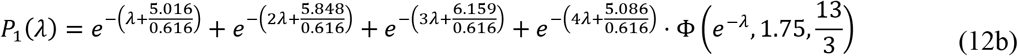

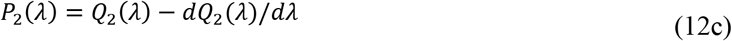

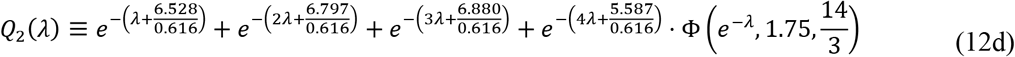

where Φ is the Lerch transcendent (22). These expressions are based on a baseline free energy of each GC|CG doublet being −3.42 kcal/mol stable (23–25). Depending on the identity of N in the (CNG) repeat unit, there are additional stabilizations resulting from the terminal mismatch, and they can be calculated using Turner’s parameters (25). For N = A, C and U, these additional bonuses are approximately ΔΔ*G* = −0.9, −0.5 and −1.6 kcal/mol per duplex, respectively. In Eq.(12a), we have added an extra term *λ*_•_ = ΔΔ*G*/*RT* to the exponent for each duplex (dot) to allow for this. In general, *λ*_•_ can be thought of as a chemical potential for the duplexes. For the case of N = G, the extra stabilization when the duplex is adjacent to a hairpin is −5.2 kcal/mol, but it is different at −7.9 when adjacent to a 2-way junction. This disparity can be accommodated by assigning different *λ*_•_ values to the numerator and denominator in the last term in Eq.(11), but this paper will not address (CGG) repeats as they can potentially also form quadruplexes {companion paper}.

Since the generating function for *Z* sums over all *n*, the value of *λ* that corresponds to the desired target value for ⟨*n*⟩ must first be identified using the relationship ⟨*n*⟩ = −∂ ln *Z* /*∂ λ*, similar to what is typically done in the grand canonical ensemble. The result is shown in Fig. 12, where the solid blue line shows ⟨*n*⟩ as a function of *λ*. Since the critical expansion threshold associated with disease onset in TREDs is *n* > 60, we employed 60 as the target value for ⟨*n*⟩. The denominator of *Z*(*λ*) changes sign at several values of *λ*, leading to divergences in *Z*. As Fig. 12 shows, these divergences are also manifested in ⟨*n*⟩. On Fig. 12, the proper value of *λ* that produces ⟨*n*⟩ = 60 is indicated by the open circle on the solid blue line, yielding *λ* = 0.1135. The other two curves in Fig. 12 relate to the effects of the stability of the duplex on the average repeat length. They will be discussed later.

**Figure 12.**
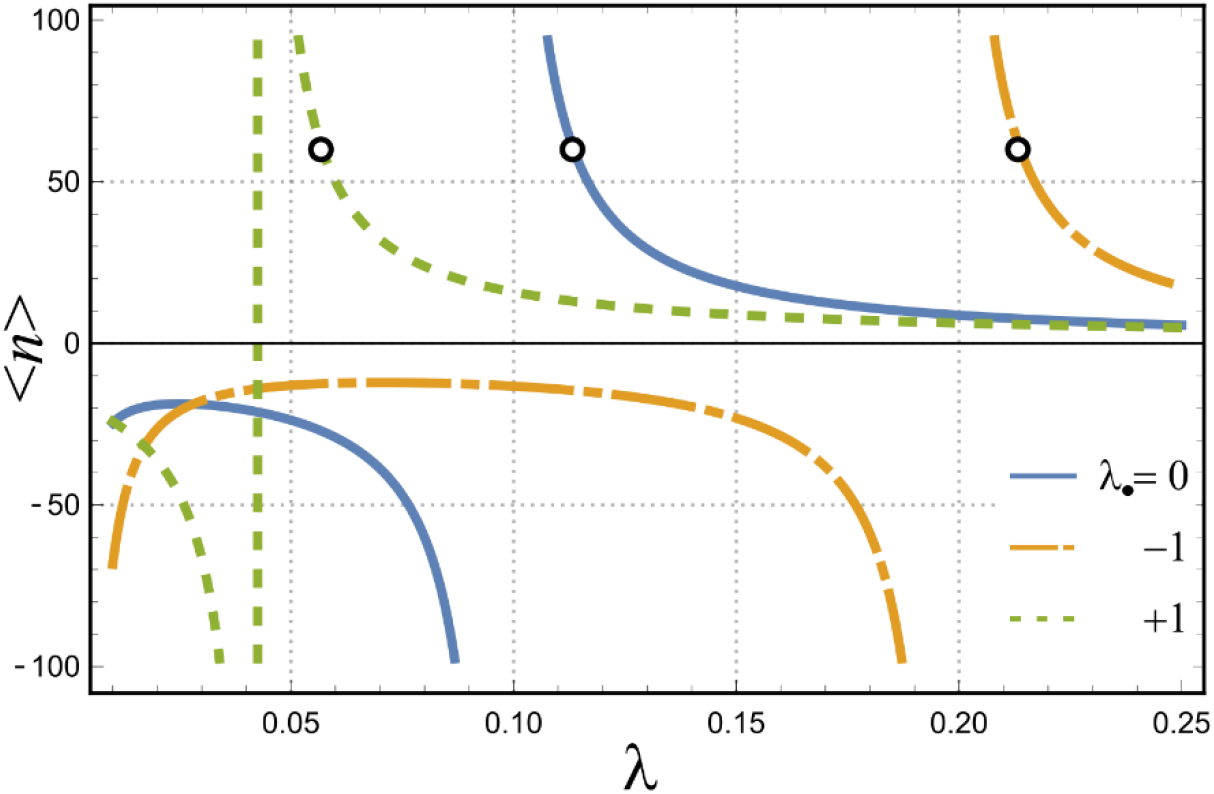
Solid blue curve shows the expectation value ⟨*n*⟩, the number of repeat units, as a function of *λ*. The point that corresponds to the target value ⟨*n*⟩ = 60 is indicated on the solid blue curve by an open circle. The dotted dash orange curve shows the same relationship when the stability of each duple is made more favorable by adding a free energy of −*RT*. The dashed green curve shows the same with an unfavorable bias of +*RT* added to each duplex.

Using the value *λ* = 0.1135, corresponding to a target ⟨*n*⟩ = 60, we calculated the expectation values for the number of duplexes, hairpins and 2-way junctions in the ensemble. These were computed by conjugating a chemical potential to each of these elements and differentiating the partition function against each of them. The results are shown in Fig. 13. The blue solid line in Fig. 13 shows the average number of repeats and is identical to the solid blue line in Fig. 12, except with only the relevant domain of *λ* displayed in Fig. 13. The open circle on the solid blue line in Fig. 13 indicates the target value of ⟨*n*⟩ = 60, and the vertical thin dashed line, the value of *λ* that produces it. The green long dashed line in Fig. 13 shows the average number of duplexes found on the sequence, giving ⟨*n*_•_⟩ ~ 4.35 for repeat length ⟨*n*⟩ = 60. The purple short dashed line in Fig. 13 shows the average number of hairpins found on the sequence, and ⟨*n*_1_⟩ ~ 4.07 for repeat length ⟨*n*⟩ = 60. The red dotted dashed line in Fig. 13 shows the average number of 2-way junctions found on the sequence, and ⟨*n*_2_⟩ ~ 0.28 for ⟨*n*⟩ = 60. Because of the specific topologies of those diagrams included in the necklace approximation of the root function, the number of hairpins plus the number of 2-way junctions must equal the number of duplexes, and this conservation law is consistent with the observed expectation values.

**Figure 13.**
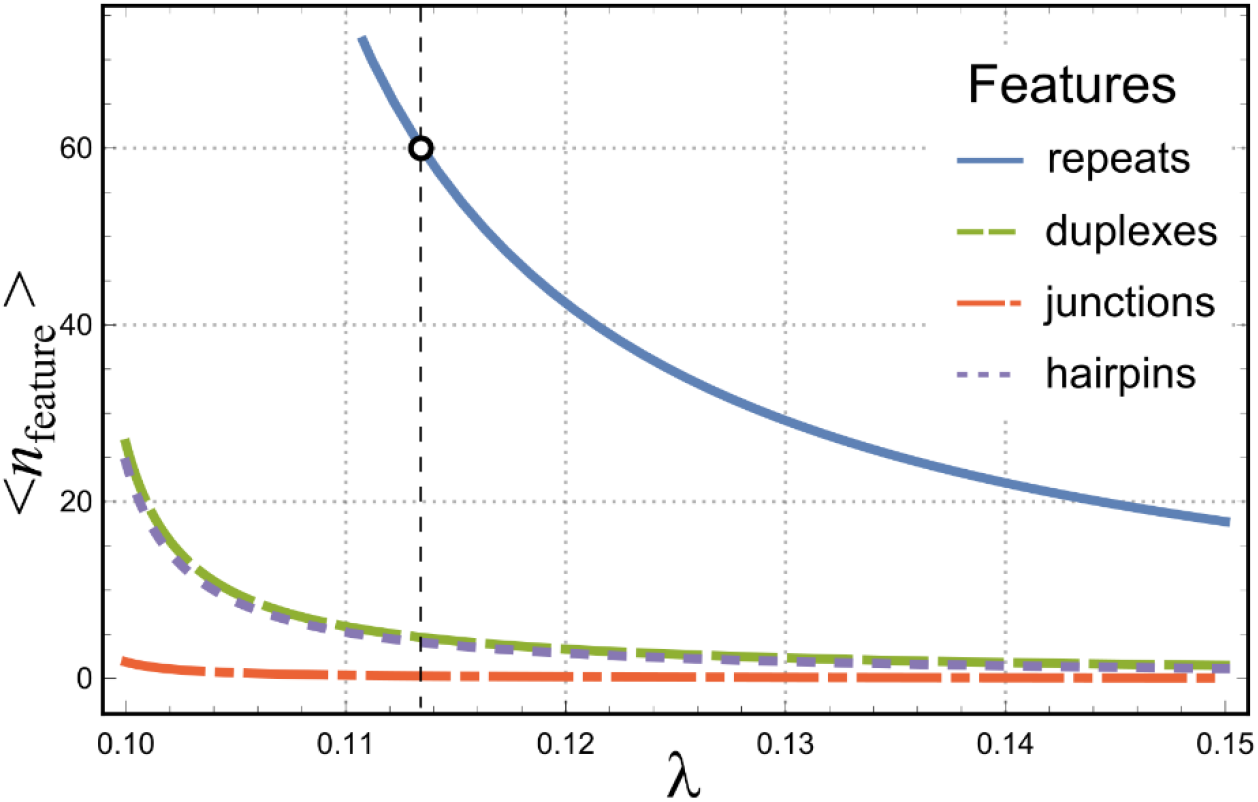
Expectation values of different features in an equilibrium ensemble of (CNG) repeat sequences at 310 K as a function of *λ*. Solid line (blue): (CNG) repeats. Long dashed line (green): duplexes. Dotted dashed line (red): 2-way junctions. Short dashed line (purple): hairpin loops. Vertical thin dashed line indicates the proper value of *λ* that produces the target value for ⟨*n*⟩ = 60, indicated by the open circle on the solid blue line.

The ensemble average of the number of duplexes, ⟨*n*_•_⟩ is a measure of the secondary structure contents on the chain at equilibrium. The maximum number of duplexes, in the case of (CNG)_n_ repeat sequences, that can be accommodated on the chain is *n*_•,max_ = ⟨*n*⟩/2. This corresponds to the case when the sequence folds into a single conformation, corresponding to the longest necklace structure possible. This “maximal” necklace conformation has been implicitly assumed to be the dominant structure for (CNG) repeat sequences at equilibrium. But the data in Fig. 13 suggest that this is not the case. For (CNG)*n* chains with *n* = 60, the dominant equilibrium conformations are instead characterized by largely open strands, with only a few duplexes, ~4 to 5. From the second derivative of *Z* with respect to the chemical potential of the duplexes, we also computed the variance in the number of duplexes, 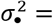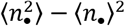, which turned out to be ~ 24.0. This suggests that while the average number of duplexes on the chain is ~4 to 5, the equilibrium fluctuations are rather large, with a standard deviation ~ 4.9 repeat units. In order words, the equilibrium conformations of the chain in a thermal ensemble at *T* = 310 K are rather diverse, where the expectation value of the number of duplexes is ⟨*n*_•_⟩ = 4.35 ± 4.9 (CNG) units, suggesting that a rather broad distribution of secondary structural contents is expected to characterize (CNG) repeat sequences in solution. The assumption that the ensemble should be dominated by the conformation with the largest number of 2-way junctions and duplexes corresponding to the maximal-length necklace, is unfounded.

Fig. 14 shows the sensitivity of the chain’s expected duplex content on free energy bias applied to different structural elements on the chain. The three panels in Fig. 14 are similar to Fig. 12, but instead of showing ⟨*n*⟩ on the vertical axis, they show the “duplex fraction”, defined as the average number of duplexes ⟨*n*_•_⟩ divided by the number of repeats ⟨*n*⟩, plotted as a function of *λ*, the chemical potential conjugate to the number of repeats. The duplex fraction is a primary indicator of the level of secondary structure contents on the chain. The theoretical maximum for the duplex fraction is 0.5, corresponding to the maximal necklace structure. Fig. 14(a) shows the effects of adding a bias *λ*_•_ to the duplexes: the orange dotted dashed line for a favorable bias *λ*_•_ = −1, the green dashed line for an unfavorable bias *λ*_•_ = +1 and the solid blue line for no bias. Similarly, Fig. 14(b) shows effects of adding a bias to the 2-way junctions, *λ*_2_ (favorable bias *λ*_2_ = −1 in the orange dotted dashed line, unfavorable bias *λ*_2_ = +1 in the dashed green line, no bias in the solid blue line). Fig. 14(c) shows the same, but for a bias applied to the hairpins (*λ*_1_ = −1 for orange dotted dashed line, −1 for green dashed line and 0 for solid blue line.) And as a point of reference, the solid blue lines in all three panels are the same curve, since they correspond to 0 bias on all three elements. In each panel, the value of the duplex fraction corresponding to the proper target length ⟨*n*⟩ = 60 under each set of biases are identified by circles. These were found by using Fig. 12, in a procedure similar to what has described above. The solid blue line in Fig. 12 describes how ⟨*n*⟩ depends on *λ*, and the orange dotted dashed line shows the same in the presence of a favorable biased *λ*_•_ = −1 placed on the duplexes, whereas the bias *λ*_•_ = −1 is unfavorable in the green dashed line. The circles in Figl. 12 show the proper values of *λ* corresponding to the target length ⟨*n*⟩ = 60. These values are also highlighted by the circles in Fig. 14(a). A similar procedure was used to identify the proper values of *λ* when biases are placed on either the junctions or the hairpins, but leaving out the details, these are also highlighted by the circles in Figs. 14(b) and (c).

**Figure 14.**
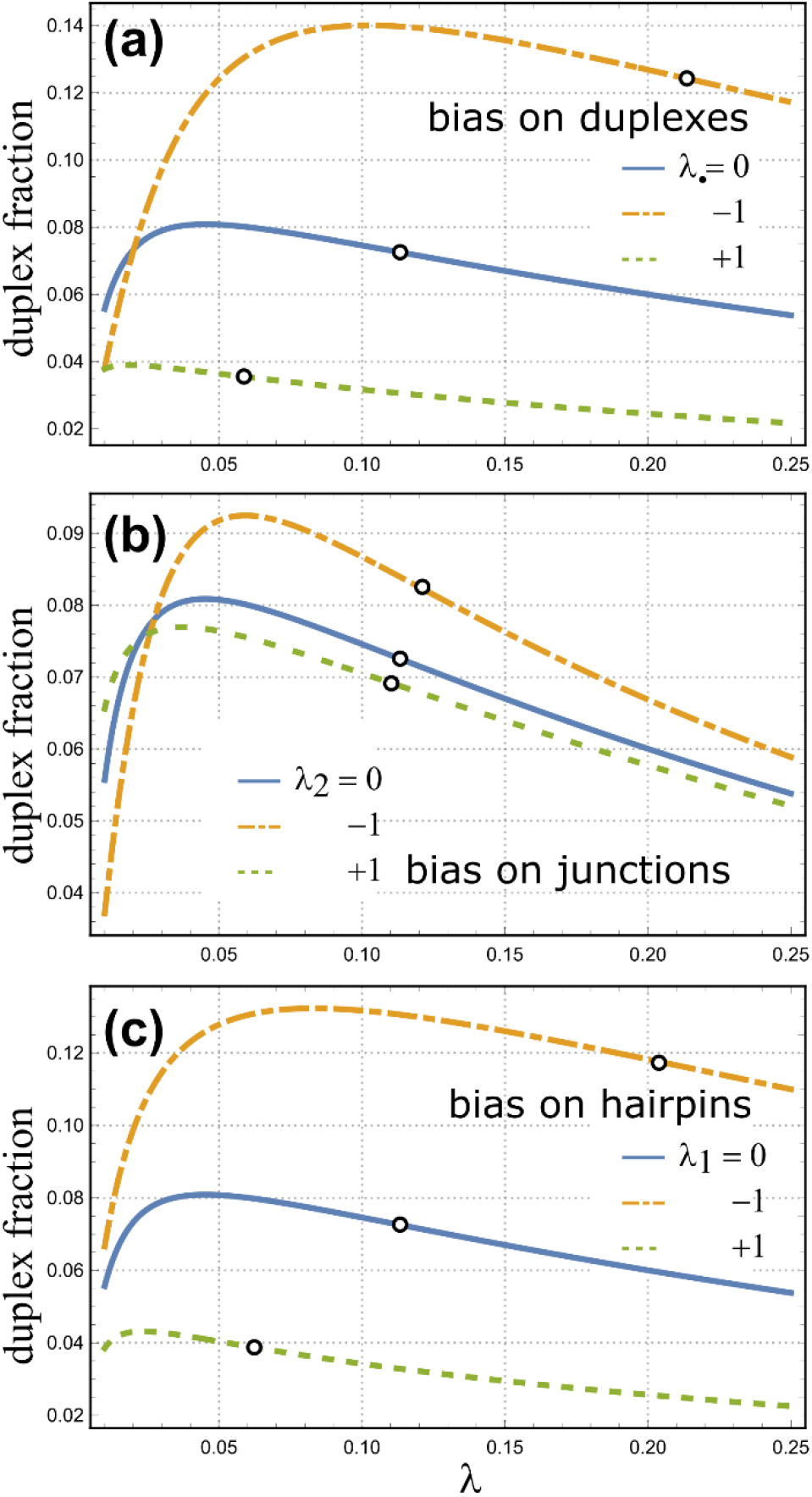
Number of duplexes per repeat units as a function of λ when perturbations are applied to bias the stability of different elements on the chain. (a) Bias applied to duplexes. (b) Bias applied to 2-way junctions. (c) Bias applied to hairpin loops. In all three panels, solid blue line indicates no bias. Dotted dashed orange line indicates a favorable bias of −*RT* applied to the free energy of each element. Dashed green line indicates an unfavorable bias of +*RT* applied. For each curve, the proper value of λ needed to produce the target ⟨*n*⟩ =60 is indicated by an open circle. The solid blue lines in all three panels are the same curve, since they correspond to 0 bias on all three elements.

Fig. 14 demonstrates that no matter whether the bias is applied to the duplexes (*λ*_•_ in (a)), 2-way junctions (*λ*_2_ in (b)) or the hairpins (*λ*_1_ in (c)), it influences the formation of duplexes in the same direction, i.e. favorable biases on either duplexes, junctions or hairpins produced more duplexes, and unfavorable biases on any of these three produced fewer duplexes. This is not surprising, because the formation of hairpins, junctions and duplexes are correlated with each other. Biasing any one of these elements will have an overall effect on the number of duplexes, which is the primary indicator of the level of secondary structure contents on the chain. However, it is also clear that biasing the duplexes has the largest impact, whereas biasing the 2-way junctions has the least. (Note difference in vertical scales among the three panels.) Compared to the effects resulting from biasing the duplexes or biasing the 2-way junctions, the impact of biasing the hairpins on the chain’s secondary structures is slightly weaker than the effect of biasing duplexes, but a bit stronger than that coming from biasing junctions.

Here, we want to make one interesting observation on the graphs in Fig. 14. The same domain of *λ* values used for Fig. 12 is also used for Fig. 14. But while Fig. 12 shows there are divergences in the expectation value of ⟨*n*⟩ as a function of *λ*, there are no apparent divergences in the duplex fraction in Fig. 14, after the number of duplexes ⟨*n*_•_⟩ *has been divided by* ⟨*n*⟩. This demonstrates an interesting feature in diagrammatic resummation, where the radius of convergence in many of the infinite sums can be effectively extended in the complex plane around singularities when certain thermodynamic observables are considered. For example, in this case of the duplex fraction, there is are discontinuities in ⟨*n*_•_⟩/⟨*n*⟩ even at the places where the denominator of the partition function *Z*(*λ*) changes sign.

The data in Fig. 14 characterizing how secondary structure contents on the chain respond to biases can also be understood in terms of the susceptibility of ⟨*n*_•_⟩ to variations in the chemical potentials of duplexes, *λ*_•_, junctions, *λ*_2_ and hairpins, *λ*_1_. For example, the susceptibility *χ*_••_ ≡ ⟨∂*n*_•_/∂*λ*_•_⟩_*λ*_•_=0_ can be computed directly from the partition function *Z*(*λ*), giving *χ*_••_ = −23.9. The other susceptibilities for the effects of junctions and hairpins on the formation of duplexes can be computed analogously, giving χ_•2_ = −1.82 and χ_•1_ = −22.1, respectively. Similarly, the susceptibilities of junctions and hairpins can also be calculated the same way, yielding χ_2•_ = χ_•2_ = −1.82, χ_22_ = −0.399, χ_21_ = −1.43, χ_1•_ = χ_•1_ = −22.1, χ_12_ = χ_21_ = −1.43 and χ_11_ = −20.6. These susceptibilities agree with the directions and magnitudes of the effects observed in Fig. 14.

In Fig. 15, we show how the duplex fraction varies as a function of different biases, in units of *RT*, imposed on the duplexes *λ*_•_ (solid lines) or on the hairpins *λ*_1_ (dashed line). Whereas in Fig. 14, the applied biases are either ±1, Fig. 15 shows the duplex fraction as a continuous function of the bias on the duplexes *λ*_•_ or the hairpins *λ*_1_, for ⟨*n*⟩ fixed at 60. When the bias is applied to the duplexes, i.e. when *λ*_•_ is varied, there are actually two families of solutions. These are shown in Fig. 15 as the orange solid line, labeled the “open” branch, and the cyan solid line, labeled the “necklace” branch. In contrast, there is only one physical solution when the bias is applied to the hairpins, i.e. when *λ*_1_ is varied, and this is indicated by the dashed line in Fig. 15.

**Figure 15.**
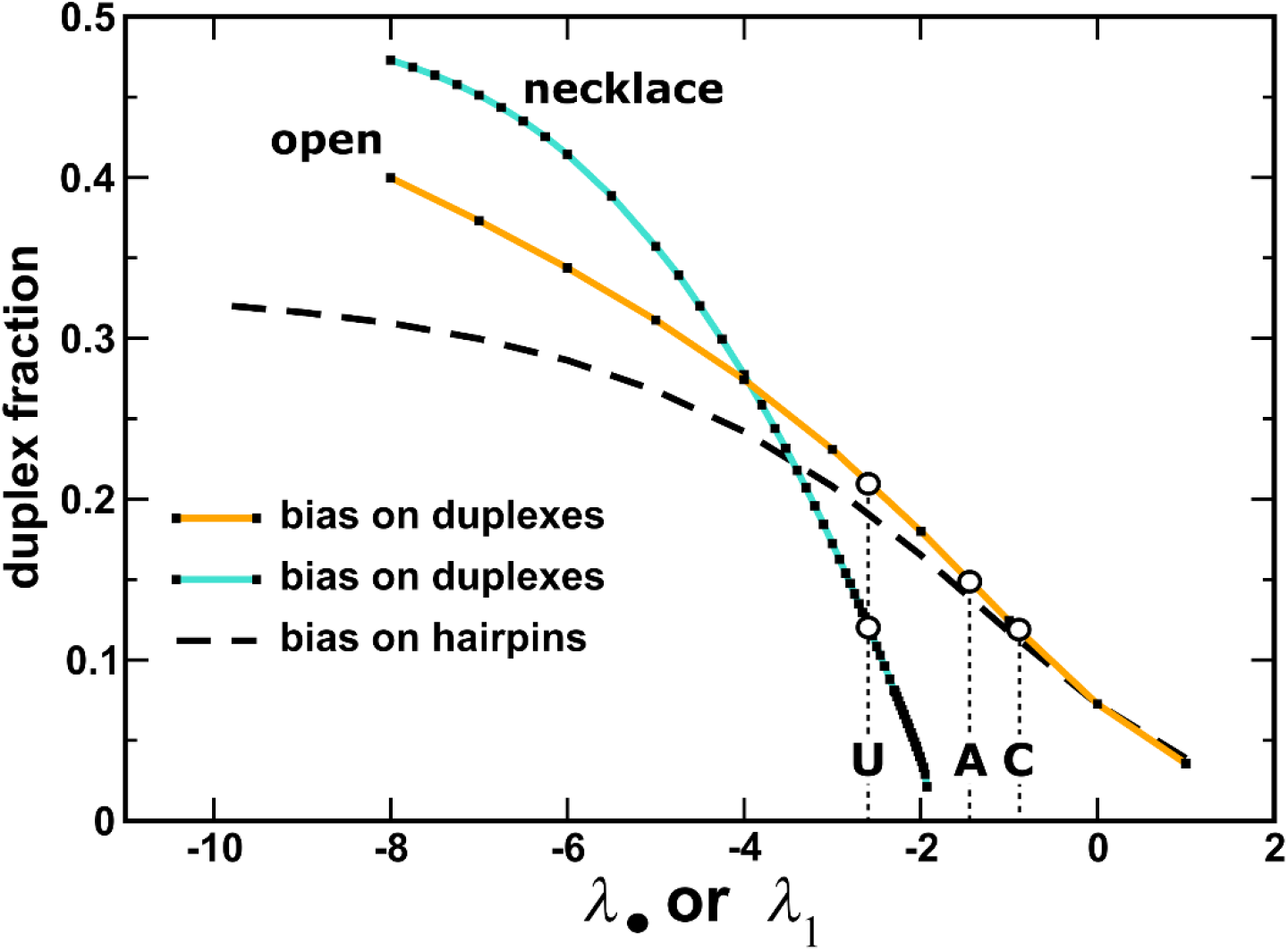
Duplex fraction as a function of bias applied to either the duplexes, *λ*_•_, or the hairpins, *λ*_1_. Theoretical maximum for the duplex fraction is 0.5. Circles indicate duplex fractions with *λ*_•_ biases associated with N = C, A or U.

Up to this point, we have only been discussing solutions on the “open” branch. The data presented in Fig. 12, 13 and 14 above are all associated with *λ*_•_ = 0 where there is only one physical solution on the “open” branch. And as we have described above, these solutions on the “open” branch are characterized by largely open chains with a moderate duplex fraction that varies as a function of the bias applied to the duplexes. The majority of the chain’s secondary structures on the “open” branch are hairpins, with a very minor fraction in the form of 2-way junctions.

As we have described under Eq. (12) above, depending on whether N = A, C or U, there are additional free energy bonuses associated with the duplexes, ΔΔ*G* = −0.9, −0.5 and −1.6 kcal/mol per duplex, respectively. The values of the *λ*_•_ bias corresponding to these values are indicated in Fig. 15 by the vertical dashed lines, and the corresponding solutions for the duplex fraction highlighted by open circles. Fig. 15 shows that different (CNG) repeat sequences with N = C, A or U are expected to produce somewhat different duplex contents on the chain. Their structural ensembles, however, are physically similar since they are all characterized by solutions on the “open” branch, i.e. they have large structural diversities as well as large equilibrium fluctuations.

Before discussing the second branch, we examine the dashed line in Fig. 15, which describes how the duplex fraction varies as a function of bias imposed on the hairpins *λ*_1_. These results were obtained by varying *λ*_1_, while keeping *λ*_•_ = 0, i.e. no additional bias on the duplexes. These solutions as a function of *λ*_1_ belong to the same physical family of solutions as the “open” branch as a function of *λ*_•_, so we can compare them with each other. For small biases for either *λ*_•_ or *λ*_1_ ≥ −3, the effect is only slightly stronger when the bias is applied directly to the duplexes, compared to when the bias is applied to the hairpins. This may seem somewhat puzzling because one would expect a perturbation applied directly to the duplexes should have a stronger effect on the duplexes, but this result is nonetheless consistent with results from Fig. 14. Considering the diagram that are included in the root function (see Fig. 10), it is clear that one duplex is always paired with a hairpin, while at the same time, one duplex is also paired with every 2-way junction. The fact that the effects on duplex formation are similar regardless whether the bias is imposed directly on the duplexes or on the hairpins implies that the number of 2-way junctions must be small in the ensemble, and Fig. 13 also corroborates this. When the magnitude of the bias becomes larger beyond *λ*_•_ or *λ*_1_ < −3, the difference between them becomes more significant. In this region, direct perturbations applied to the duplexes do initiate more duplexes in comparison to when the bias is applied to hairpins. The duplex fraction, in the limit of very large bias imposed on hairpins, appears to approach an asymptotic value of ~ 0.33, whereas it approaches the theoretical maximum of 0.5 when the bias is applied directly to the duplexes. The bifurcation between these two curves suggests that the number of 2-way junctions in the ensemble becomes more significant when a larger bias is placed on the duplexes directly.

We now turn to the “necklace” branch. This family of solutions are shown as the cyan solid line labeled “necklace” in Fig. 15. As the bias applied to the duplexes *λ*_•_ is increased, a second and physically distinct family of solutions appears. This new branch of solutions is characterized by a duplex fraction that grows in rapidly as a function of *λ*_•_. This branch appears abruptly when *λ*_•_ reaches the critical value ~ − 1.9, where the duplex fraction on the cyan solid line begins to take on a nonzero value. Once the second branch is seeded, its duplex fraction increases rapidly toward its theoretical maximum value of 0.5. We have also computed the expectation values of hairpins and 2-way junctions along this so-called “necklace” branch, and the average number of hairpins is close to one along the entire branch. The ensembles characterized by solutions on this new branch are therefore made up of the type of necklace structures illustrated in Figs. 1, 3 and 4, with a stretch of successive two-way junctions interposed by shorts helixes and with a single hairpin cap. This is not surprising, because given sufficiently stable duplexes, one would expect the chains to want to maximize base pairing even at the cost of lowering conformational entropy and diversity, in order to fold into these necklace structures with increasing number of junctions, until they grow to maximum length, at which point the duplex fraction becomes 0.5. Fig. 15 shows that this is indeed the asymptotic behavior displayed by this branch of solutions. But while this suggests that the necklace diagrams in Figs. 1, 3 and 4 are indeed predicted by the diagrammatic theory here, we note that the magnitude of the bias needed to be applied to the duplexes are quite large for these necklace solutions to become physically relevant.

Fig. 15 shows that for N = C or A, (CNG) repeat sequences only have a single solution on the “open” branch. But for N = U, the stronger stability of duplexes with a U:U terminal mismatch generates a second solution on the “necklace” branch also, and because of this, (CUG)_n_ repeat sequence could produce physically distinct ensembles compared to (CAG)_n_ or (CCG)_n_ chains. But while this is true, Fig. 15 also shows that at the value of *λ*_•_ ~− 1.6 for N = U, the duplex fraction on the “necklace” branch is actually lower than on the “open” branch. This suggests that while it is possible for (CUG)_n_ chains to develop an alternative subensemble dominated by necklace structures, there is actually an overall *higher* level of secondary structure contents in the “open” ensemble. Because of this, we can conclude that for all three N = C, A or U, the equilibrium conformations of long (CNG)_n_ repeats are dominated by a low level of secondary structures at 310 K.

## CONCLUSION

We have formulated a diagrammatic theory to study the conformational ensembles of (CNG)_n_ RNA sequences. Transcripts of overexpanded microsatellites on the genome containing 60 to 100 (CNG) repeats have been implicated in a number of neurological diseases known as TREDs. To understand the structures of these (CNG) repeats sequences, we performed a series of calculations aimed at characterizing their equilibrium ensembles. Based on a diagrammatic representation of the partition function, our calculations are based on using graphs to annotate secondary structural motifs on the chains, and, in conjunction with evidence from previous simulation studies, these diagrammatic representations allowed us to easily factorize the graphs in order to re-express the free energy of each configuration as a sum of independent terms. Using generating function mathematics and diagrammatic resummation techniques, we were able to derive a closed-form expression for the partition function in terms of a renormalized root function, which is the diagrammatic equivalence of the sum over all self-contained circuit diagrams. Employing a simple approximation for this root function, we derived analytical expressions for the partition function and its corresponding thermodynamic observables. With the so-called necklace approximation for the root function, the partition function captures an infinite set of conformations with any number and any combination of duplexes, hairpins and 2-way junctions. Together with simulation data from a self-consistent library of thermodynamic costs previously obtained for the various graph elements, we solved the resulting equations to arrive at numerical estimates for the ensemble expectation values of duplexes, hairpins and 2-way junction on the chain, enabling us to quantitatively characterize the structural diversity of different (CNG)_n_ ensembles.

While most studies in the field have implicitly assumed that the ensemble of a (CNG)_n_ sequence is dominated by a single structure having the maximal number of paired bases forming duplexes interposed by 2-way junctions between them, the results of this study suggest otherwise. The data show that the structural ensembles of (CNG)_n_ repeat sequence with n ~ 60 are surprisingly diverse. The equilibrium number of duplexes, hairpins and junctions on these sequences indicate that their secondary structure contents are far from the expected maximally paired conformation. To the contrary, the ensemble is dominated by open segments, with approximately 8 to 13 duplexes on the average from (CCG)_n_ to (CUG)_n_ chains, whereas the maximally paired structure would have 30 duplexes. But given the exaggerated equilibrium fluctuations on the structure, the number of duplexes could be as high as 16 to 23. The results show that the majority of the loops are in the form of hairpins, but the variance in the secondary-structural contents on the chains are also quite large within these ensembles. And due to these large equilibrium fluctuations, the chains are also highly suspectable to external perturbations. The results show how perturbations in the form of biases on the stabilities of the various structural motifs, duplexes, junctions and hairpins, could affect the secondary structures of the chains in either directions. It appears that the number of duplexes and hairpins are also strongly correlated even though the overall level of secondary structure exhibit strong equilibrium fluctuations, and because of this, it is quite possible that the secondary structures could be easily pinned by tertiary contacts between different parts of the chain, either from the chain interacting with itself or from interactions mediated by the binding or association of the chain to ligand(s). This may in turn have implications on how these (CNG)_n_ sequences could acquire unintended functions in the cell, leading to their cytotoxicity. Studies are underway to examine whether these perturbations could indeed quench the structural fluctuations of the chains. We are also investigating different classes of approximations for the root function to try to ascertain whether other structural elements, such as multiway junctions or pseudoknots, could also play important roles in controlling the structural ensembles of these (CNG) repeat sequences.

## AUTHOR CONTRIBUTIONS

CHM designed the study. CHM and ENHP carried out the work. CHM and ENHP wrote the manuscript.

## ACKNOWLEDGEMENT

This material is based in part upon work supported by the National Science Foundation under Grant Number CHE-1664801.

